# Feeding the disparities: the geography and trends of breastfeeding in the United States

**DOI:** 10.1101/451435

**Authors:** Romain Garnier, Ana I. Bento, Pejman Rohani, Saad B. Omer, Shweta Bansal

## Abstract

There is scientific consensus on the importance of breastfeeding for the present and future health of newborns, in high- and low-income settings alike. In the United States, improving breast milk access is a public health priority but analysis of secular trends are largely lacking. Here, we used data from the National Immunization Survey of the CDC, collected between 2003 and 2016, to illustrate the temporal trends and the spatial heterogeneity in breastfeeding. We also considered the effect sizes of two key determinants of breastfeeding rates. We show that, while access to breast milk both at birth and at 6 months old has steadily increased over the past decade, large spatial disparities still remain at the state level. We also find that, since 2009, the proportion of households below the poverty level has become the strongest predictor of breastfeeding rates. We argue that, because variations in breastfeeding rates are associated with socio-economic factors, public health policies advocating for breastfeeding are still needed in particular in underserved communities. This is key to reducing longer term health disparities in the U.S., and more generally in high-income countries.

## Introduction

Maternal breast milk represents the best source of nutrition for newborns, boosting the development of brain function, and providing short and long-term benefits to individual health (UNICEF & WHO, 2018). In addition, breastfeeding leads to the transfer of an array of immune compounds (IgA and to a lesser extent IgG) from mother to infant allowing specific protection against respiratory and enteric pathogens during a time when the risk of severe outcomes is highest (Victora et al., 2016), and shaping the establishment of the infant’s gut microbiota (Pannaraj et al., 2017). The important role of breastmilk in early life health is illustrated by the half-fold infectious disease mortality among breastfed infants (compared to non-breastfed) in low and middle income countries (Victora et al., 2016), and by the 53% reduction in monthly diarrheal hospital admissions due to exclusive breastfeeding in a high income country (Quigley, Kelly, & Sacker, 2007). Because of the nutritional, immunological, and developmental benefits, hundreds of thousands of annual deaths in children under five years of age are expected to be prevented by global improvements in access to breastfeeding (UNICEF & WHO, 2018; Victora et al., 2016), and high income countries would also receive potentially large economic benefits (Rollins et al., 2016). The scientific consensus around the importance of breastfeeding has led the World Health Organization (WHO) to set a target of 50% of infants under 6 months old being exclusively breastfed. However, this target remains elusive, and concerns with access to breastfeeding has prompted the World Health Assembly, during its 2018 meeting, to issue a resolution in support of the practice (Seventy-first World Health Assembly, 2018). The resolution emphasized the need for further efforts in supporting continued breastfeeding, in particular in high-income countries.

In the United States, the Healthy People 2020 program highlights breastfeeding as a public health priority and sets targets of breastfeeding initiation among 81.9% of births and of continuation of breastfeeding until six months of age in 60.6% of infants by year 2020. Suboptimal breastfeeding in the United States is responsible for additional maternal and infant deaths, and carries economic costs in the order of the billions of dollars (Bartick et al., 2017). Achieving the Healthy People 2020 goals will require ensuring socio-demographic equity in breastfeeding rates. A previous study showed that black infants were generally less likely to be breastfed, and that the magnitude of this difference varied extensively depending on the state (Anstey, Chen, Elam-Evans, & Perrine, 2017). However, comprehensive analyses of recent secular trends and spatial heterogeneity in breastfeeding rates in the United States are still lacking.

## Methods

Annual data on breastfeeding is available from the Center for Diseases Control and Prevention (CDC) for the 2003-2016 period, as part of successive rounds of the National Immunization Survey (NIS). The NIS represents a unique long-term source of comparable data at the state level and has so far been largely underused for analyses of spatial and temporal breastfeeding trends. Data is directly available on the rates of initiation (“breastfeeding initiation”). Data on the rates of breastfeeding at 6 months old (“breastfeeding continuation”) are derived from information collected on the duration of breastfeeding.

We analyzed trends of breastfeeding using Mann-Kendall tests, and tested the effect of the proportion of black people and of households below the poverty level using multiple linear regressions. All statistical analyses were run in R 3.5.0.

## Results

Breastfeeding initiation in the United States follows a significant positive trend over the past 14 years (Mann-Kendall test; p < 0.001) to reach a maximum of 84.4% in 2016 (Figure 1). Similar significant positive trends are found in all but four states (Arizona, Georgia, Montana, and Utah; no significant trend in these states) indicating that the upwards trend is robust across the country.

**Figure 1.**
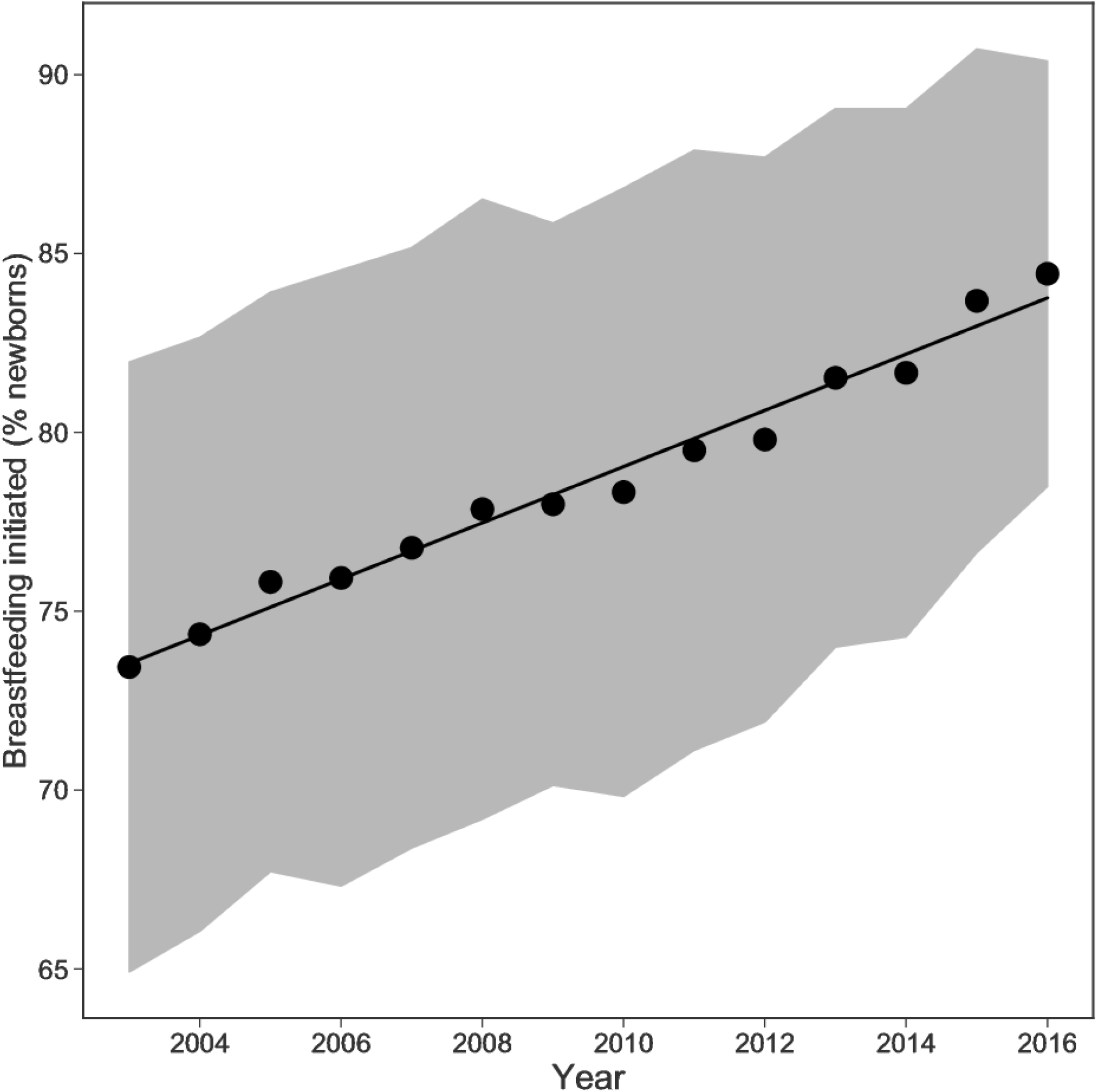
Breastfeeding rates in the United States between 2003 and 2016, estimated from the National Immunization Survey. The significant trend line is estimated from a linear model and the shaded area corresponds to one standard deviation around the mean.

Breastfeeding rates at 6 months are highly correlated with rates of initiation (R2 = 0.75; p < 0.001; Figure S1) and follow a significant positive trend at the national scale (Mann-Kendall test; p < 0.001), reaching 58.6% in 2016. This upward trend in continuation of breastfeeding to 6 months is also found in all but five states (Alabama, Kentucky, New Hampshire, New Mexico, and Oregon). We note that there is no overlap between these states where increase in continuation rates was not significant and the states where the increase in initiation rates was not significant.

Despite the overall national increase, substantial spatial variation in breastfeeding rates persists in 2016 (Figure 2), varying between 65.3% in Mississippi at the lowest and 93.4% at the highest in Oregon. In addition, twenty states had a breastfeeding rate below the national average. Among these, the six states (Alabama, Arkansas, Kentucky, Louisiana, Mississippi, and West Virginia) with the lowest relative ratios of breastfeeding belonged to the U.S. Census South region. New policies aimed at increasing access to breast milk may thus be most effective in this region. We also found similar spatial variation in the rates of breastfeeding at 6 months (Figure S2).

**Figure 2.**
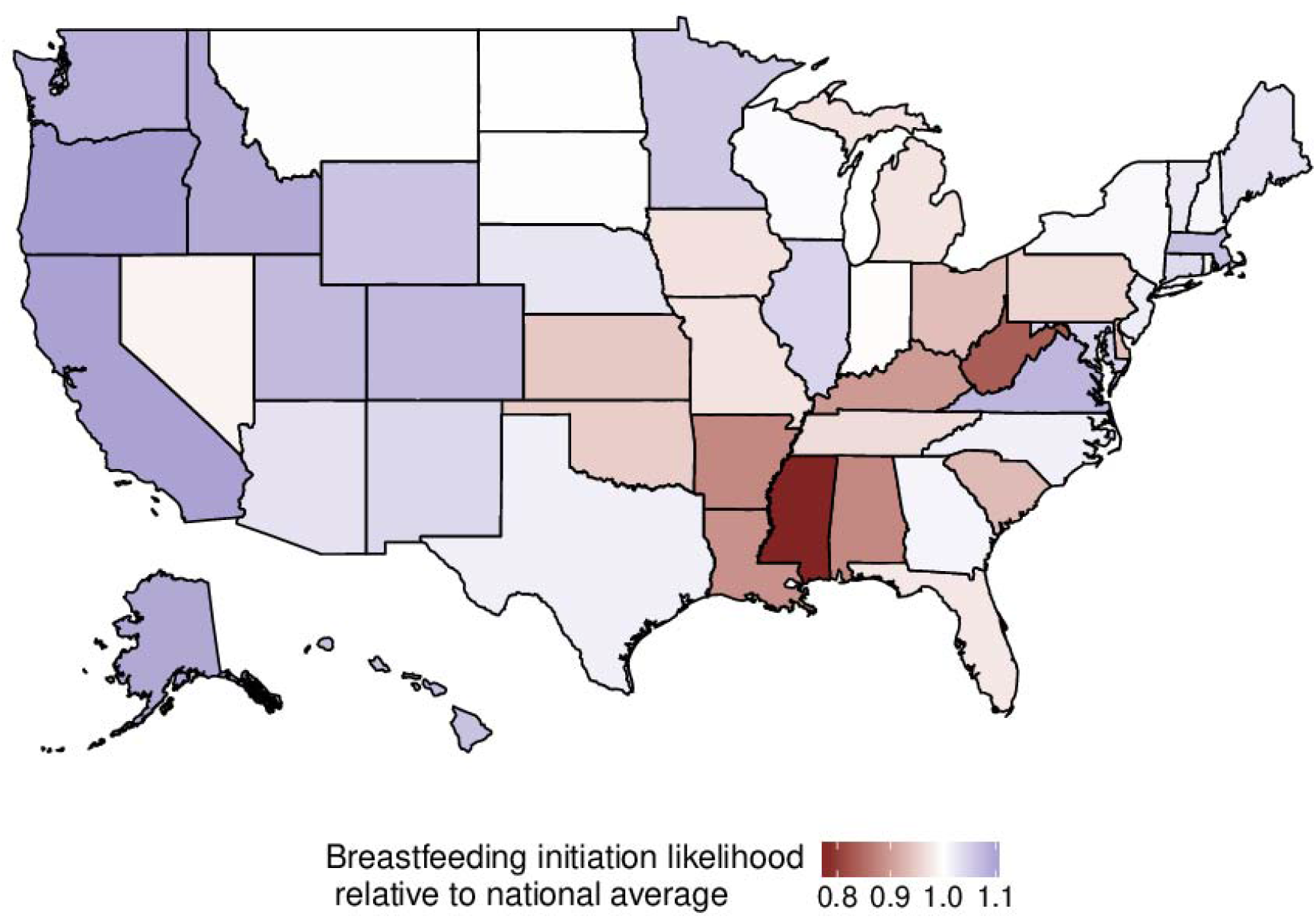
Map of breastfeeding initiation likelihood relative to the national mean (μ) for the year of 2016. A value of 1 indicates a breastfeeding rate matching μ and a value < 1 represents low breastfeeding rates relative to μ.

In 2016, we found that low levels of breastfeeding were associated with two socio-economic predictors: larger proportions of households below the poverty level (multiple linear regression: p < 0.001), and larger proportions of black infants (multiple linear regression: p = 0.02) in a state. As an illustration, we separated states with an above-average rate of breastfeeding (mean = 88.2%) from states below the national average (mean = 79.9%). Similar to Anstey et al. (2017), we found a 4.1 percentage points increase in the average proportion of black infants in states with low breastfeeding compared to states with high breastfeeding. Further, we found an even stronger difference in the proportion of households below the poverty line: 25.4% of families were below the poverty line in states with low breastfeeding, compared to 18.1% in states with high breastfeeding. While it is difficult to fully disentangle the racial and socio-economic contexts, we nevertheless found that their respective influences varied throughout the period covered by the NIS data (Figure 3). The association with income first appeared significant in 2009 while the proportion of black infants appeared to become less predictive after that date. The association between income and breastfeeding rates at 6 months followed the same pattern (Figure S3). However, breastfeeding at 6 months appeared less often to be significantly associated with the proportion of black people in a state, in particular in the recent years covered in the NIS dataset.

**Figure 3.**
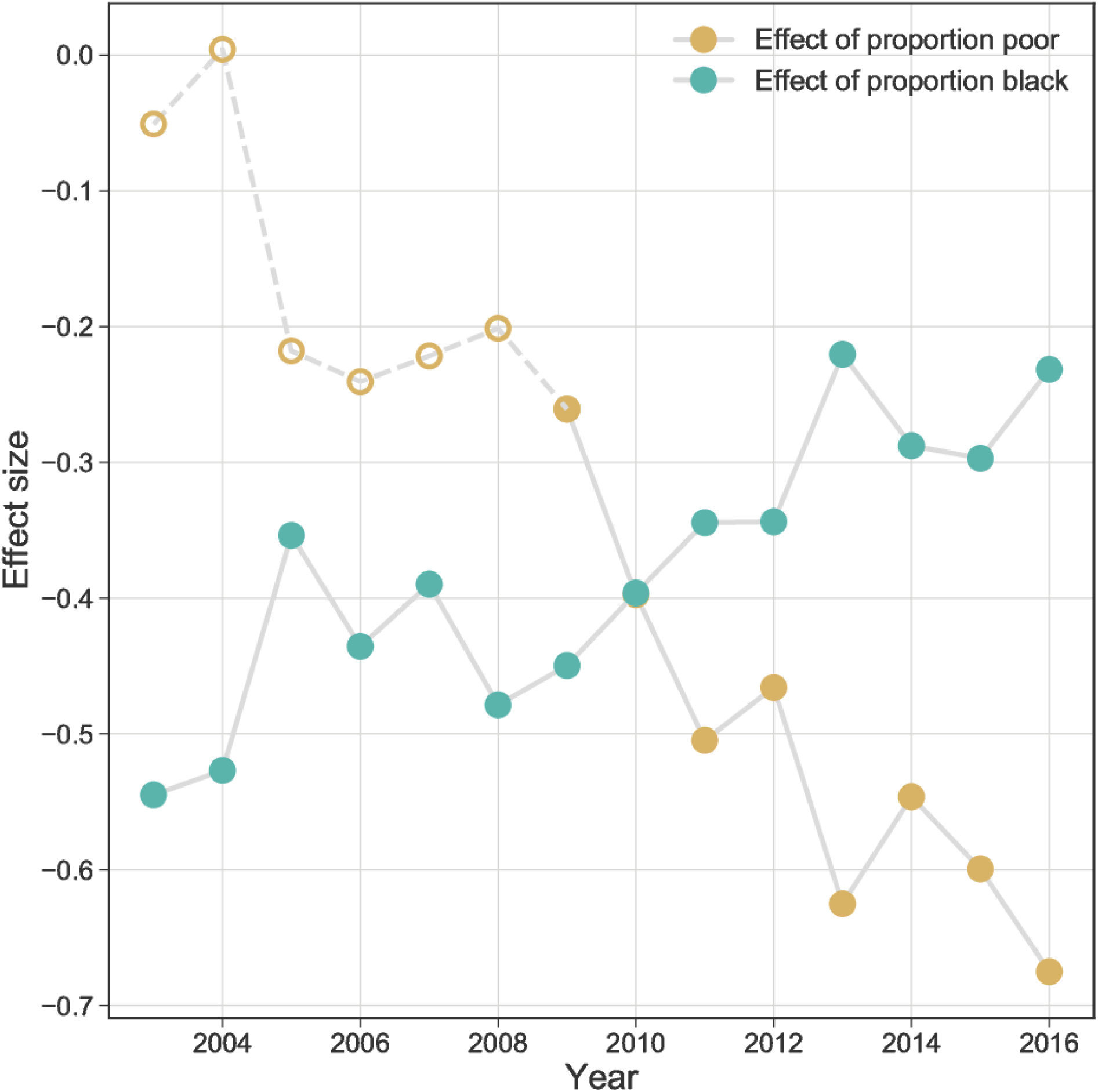
Estimates of the association of poverty (yellow) and being black (green) with the rates of breastfeeding initiation during 2003-2016. The open circles indicate non-significant estimates and closed circles indicate significant (p < 0.05) estimates.

## Discussion

Overall, the positive trend in breastfeeding initiation in the United States is in line with the targets set by public health policies. However, breastfeeding rates at six months old remain below the target set by the Healthy People 2020 program (58.6% in 2016 versus the target of 60.6%), but this target remains achievable by the end of the program. Looking past national averages also reveals deep disparities among states, highlighting the need for state-level policies to remain conducive to breastfeeding and support all aspects of maternal and infant health.

Importantly, the extent of variation in breastfeeding initiation between states has remained relatively stable throughout the period of 2003-2016 (Figure 1, shaded area), suggesting that a “one size fits all” approach for all states will not be sufficient to move the national average. Ensuring that disparities are reduced will be essential in minimizing the mortality and economic burdens of inadequate breastfeeding practices in the United States (Bartick et al., 2017).

We also found that the effect of the proportion of households below the poverty line on both rates breastfeeding initiation and continuation has increased rapidly since 2009, suggesting a possible link to the economic recession of 2008-2009. Economic disparities and the amplification of socio-economic inequalities during economic crises may thus currently be important drivers of spatial variation in breastfeeding, although further research addressing the interplay of racial and economic inequalities will be necessary. This could help identify factors explaining why racial disparities appear less predictive of maintaining breastfeeding up six months.

It is also likely that the spatial heterogeneity we observe at the state-level would be even more pronounced at smaller spatial scales such as counties or zipcodes, potentially even within states with relatively high average levels of breastfeeding. Neighborhood-level socio-economic predictors can indeed be important determinants of children health outcomes (e.g. Yaeger, Moore, Melly, & Lovasi, 2018), and similar effects may be expected for breastfeeding. Future studies are needed, especially harnessing the ever improving power of spatial statistical modeling (Lee et al., 2018), and will help determine the relevant spatial scale for the design and implementation of targeted public health actions thus remains an urgent need.

As clearly outlined in a the recent World Health Assembly resolution (Seventy-first World Health Assembly, 2018), improving access to breast milk is, and should remain, a public health priority in both developing and developed countries. Concerns that women unable to breastfeed could be stigmatized, cited by the United States as reasons to oppose the resolution (Jacobs, 2018), should not stop these efforts. In the United States, and in other high income countries, future public health efforts must focus on reducing spatial disparities in breastfeeding and underlying healthcare factors where low economic status and limited access to care can further hinder positive maternal and infant health behaviors (Anstey et al., 2017). We also advocate that future epidemiological studies should focus on determining whether clusters of low breastfeeding rates spatially covary with low maternal or infant immunization rates creating pockets of double jeopardy for vaccine-preventable childhood diseases such as measles, pertussis and rotavirus. Understanding such population-level epidemiological consequences could be an important motivation to increase breastfeeding uptake in high-income countries.

## Acknowledgements

Research reported in this publication was supported by the National Institute of General Medical Sciences of the National Institutes of Health under Award Number R01GM123007. The content is solely the responsibility of the authors and does not necessarily represent the official views of the National Institutes of Health.

